# Quartet Sampling distinguishes lack of support from conflicting support in the plant tree of life

**DOI:** 10.1101/148536

**Authors:** James B. Pease, Joseph W. Brown, Joseph F. Walker, Cody E. Hinchliff, Stephen A. Smith

**Author notes:** Manuscript received_____; revision accepted_____.

## Abstract

**Premise of the Study:** Phylogenetic support has been difficult to evaluate within the plant tree of life partly due to the difficulty of distinguishing conflicted versus poorly informed branches. As datasets continue to expand in both breadth and depth, new support measures are needed that are more efficient and informative.

**Methods:** We describe the Quartet Sampling (QS) method, a quartet-based evaluation system that synthesizes several phylogenetic and genomic analytical approaches. QS characterizes discordance in large-sparse and genome-wide datasets, overcoming issues of alignment sparsity and distinguishing strong conflict from weak support. We test QS with simulations and recent plant phylogenies inferred from variously sized datasets.

**Key Results:** QS scores demonstrate convergence with increasing replicates and are not strongly affected by branch depth. Patterns of QS support from different phylogenies leads to a coherent understanding of ancestral branches defining key disagreements, including the relationships of *Ginkgo* to cycads, magnoliids to monocots and eudicots, and mosses to liverworts. The relationships of ANA grade angiosperms, major monocot groups, bryophytes, and fern families are likely highly discordant in their evolutionary histories, rather than poorly informed. QS can also detect discordance due to introgression in phylogenomic data.

**Conclusions:** The QS method represents an efficient and effective synthesis of phylogenetic tests that offer more comprehensive and specific information on branch support than conventional measures. The QS method corroborates growing evidence that phylogenomic investigations that incorporate discordance testing are warranted to reconstruct the complex evolutionary histories surrounding in particular ANA grade angiosperms, monocots, and non-vascular plants.

## INTRODUCTION

Discordance and uncertainty have emerged as consistent features throughout the history of our evolving model of the plant tree of life (Crane, 1985; Chase et al., 1993; Palmer et al., 2004; Soltis et al., 2011; Wickett et al., 2014). Particularly strong contentions often arise at pivotal transitions in the evolution of plant life on earth, such as the development of vascular tissue (Pryer et al., 2001; Steemans et al., 2009; Banks et al., 2011), the rise of seed-bearing plants (Chase et al., 1993; Chaw et al., 1997; Bowe et al., 2000; Qiu et al., 2006; Jiao et al., 2011), and the explosive radiation of flowering plants (Crane, 1985; The Amborella Genome Project, 2013; Goremykin et al., 2015; Taylor et al., 2015; Simmons, 2016; Edwards et al., 2016). Modern phylogenomic datasets, rather than quelling these disagreements, have repeatedly shown that these phylogenetic conflicts are often the result of biological processes including incomplete lineage sorting (ILS), introgressive hybridization, and paralog duplication-loss (e.g., Zhong et al., 2013b; Wickett et al., 2014; Zwickl et al., 2014; Yang et al., 2015; Eaton et al., 2017; Pease et al., 2016b; Goulet et al., 2017; Walker et al., 2017c). Several methods have been proposed to address these issues during species tree inference (e.g., Zwickl and Hillis, 2002; Ogden and Rosenberg, 2006; Shavit Grievink et al., 2010; Anderson et al., 2012; Roure et al., 2012; Hinchliff and Roalson, 2013; Mirarab et al., 2014). However, we lack a generalized framework to quantify phylogenetic uncertainty (specifically branch support) that distinguishes branches with low information from those with multiple highly supported, but mutually exclusive, phylogenetic histories.

One of the most commonly used branch support methods has been the non-parametric bootstrap (NBS; Felsenstein, 1985) and recent variants like the rapid bootstrap (RBS; Stamatakis et al., 2008), which resample the original data with replacement assuming that aligned sites are independent and identically distributed (i.i.d.) samples that approximate the true underlying distribution (Felsenstein, 1985; Efron, 1992). In practice, the assumptions of NBS (in particular site independence) may rarely be met and can deteriorate under a variety of conditions (Felsenstein and Kishino, 1993; Hillis and Bull, 1993; Sanderson, 1995; Andrews, 2000; Alfaro et al., 2003; Cummings et al., 2003). More recently the UltraFast bootstrap approximation (UFboot) method, utilizing a likelihood-based candidate tree testing, was proposed to address speed and score interpretation issues for NBS (Minh et al. 2013; and see comparison in Simmons and Norton 2014).

The other most common branch support metric has been the Bayesian posterior probability (PP). PP scores are typically calculated from posterior distributions of trees generated using a Markov chain Monte Carlo (MCMC) sampler, and summarized using a majority-rule consensus tree (e.g., Larget and Simon, 1999; Drummond and Rambaut, 2007; Holder et al., 2008; Ronquist et al., 2012; Larget, 2013). The interpretation of PP values is more straightforward than bootstrap proportions, as PP values represent the probability that a clade exists in the underlying tree, conditioned on the model of evolution employed and the prior probabilities. The individual and relative performance of PP has been well-documented as generally favorable (Wilcox et al., 2002; Alfaro et al., 2003; Cummings et al., 2003; Huelsenbeck and Rannala, 2004). However, PP may be excessively high in certain scenarios (e.g., oversimplified substitution models; Suzuki et al., 2002; Douady et al., 2003; Nylander et al., 2004). PP also may fail under a multi-species coalescent framework with conflicting phylogenies (Reid et al., 2013). This is particularly noteworthy in light of studies showing the disproportionate effects of a few genes on overall genome-wide phylogenies (Brown and Thomson, 2017; Shen et al., 2017; Walker et al., 2017a).

Ongoing efforts to expand genetic sampling to as many plant species as possible have produced increasingly species-rich, but data-sparse, alignments (i.e., large-sparse or “fenestrated” matrices). Meanwhile, the accelerating accretion of new genomes and transcriptomes will continue to deepen genome-wide datasets with millions of aligned sites. Both axes of dataset expansion present challenges to the tractability and interpretation of phylogenetic branch-support analytics. NBS scores are known to perform poorly for large-sparse matrices (Driskell et al., 2004; Wiens and Morrill, 2011; Smith et al., 2011; Roure et al., 2012; Hinchliff and Roalson, 2013; Hinchliff and Smith, 2014b), where the sampling procedure generates uninformative pseudo-replicates that mostly omit informative sites (or consist of mostly missing data). Furthermore, resampling methods (including NBS) approximate the resampling of a larger idealized population. Genomic datasets contain virtually all available data, and therefore are not samples of any larger whole. PPs provide an appropriate testing framework and straightforward interpretation for genomic data, but available Bayesian methods of analysis are not scalable to genome-wide data under current computational speeds. PPs also may over-estimate support when models are overly simple, which becomes increasingly problematic as the size and complex of datasets expand. PP and NBS scores therefore both appear unsuitable for use on large datasets, the former due to feasibility and the latter due to its assumptions (also discussed in Smith et al. 2009, Hinchliff and Smith 2014b).

As phylogenomics has developed over the last decade, alternative methods have been introduced to factor the increased data and inherent gene tree-species tree conflict. These methods measure the concordance of gene trees (broadly referring to a phylogeny from any sub-sampled genomic region), including the internode certainty (IC) and tree certainty (TC) scores (Rokas et al., 2003; Salichos et al., 2014; Kobert et al., 2016; Zhou et al., 2017), Bayesian concordance factors (Ané et al., 2006), and other concordance measures (Allman et al., 2017). These scores were developed around the central concept of a branch support statistic that measures concordance of various trees with a particular tree hypothesis. This perspective offers much for partitioning phylogenetic discordance and analyzing larger alignments more rapidly in a phylogenomic coalescent-based framework. Unfortunately, though relevant to genomic datasets, they may not be as suitable for large-sparse alignments.

Finally, quartet methods—in particular quartet puzzling methods—have been developed for phylogenetic reconstruction (Strimmer et al., 1997; Strimmer and von Haeseler, 1997; Ranwez and Gascuel, 2001; Allman and Rhodes, 2004; Chifman and Kubatko, 2014; Mirarab et al., 2014; Zwickl et al., 2014) and support (e.g., “reliability values”; Strimmer et al., 1997; Strimmer and von Haeseler, 1997). More recently, quartet procedures have been explored to facilitate sampling of large-sparse alignments (Misof et al., 2013) and as part of coalescent-based quartet inference methods (Stenz et al., 2015; Gaither and Kubatko, 2016; Sayyari and Mirarab, 2016). These quartet methods benefit from the speed advantages of a smaller alignments and the statistical consistency of quartet trees, which avoid complex lineage sorting issues that occur with more speciose phylogenies (Rosenberg, 2002; Degnan and Salter, 2005).

Despite the wide array of approaches to branch support quantification briefly discussed above, few measures (excepting concordance methods) accommodate multiple histories and distinguish different causes of poor support for a branch in the phylogeny (e.g., multiple supported-but-conflicting phylogenetic relationships vs. low information). Being able to identify a branch as having a strong consensus and a strongly supported secondary evolutionary history would provide valuable insight into the plant tree of life (among many other groups; see also Brown and Lemmon, 2007).

Here, we describe the Quartet Sampling (QS) method (summarized in Fig. 1 and Table 1), which blends aspects of many of the methods described above and leverages the efficiency of quartet-based evaluation. The goal of the QS method is to dissect phylogenetic discordance and distinguish among lack of support due to (1) low information (as in NBS and PP), (2) discordance as a result of lineage sorting or introgression (as in concordance measures), and (3) misplaced or erroneous taxa (a.k.a. “rogue taxa”; Wilkinson, 1996; Aberer et al., 2012). In many modern phylogenetic and particularly phylogenomic studies, these causes of discordance are frequently surveyed and reported separately (e.g., Xi et al., 2014a; Wickett et al., 2014; Yang et al., 2015; Pease et al., 2016b; Walker et al., 2017c). QS provides a unified method for their execution, interpretation, and reporting. Additionally, the QS method offers a viable means to describe branch support in large phylogenies built from sparse alignments (10,000–30,000 tips with >80% missing data), which are generally intractable for Bayesian analysis (though see tools like ExaBayes; Aberer et al., 2014).

**Fig. 1.**
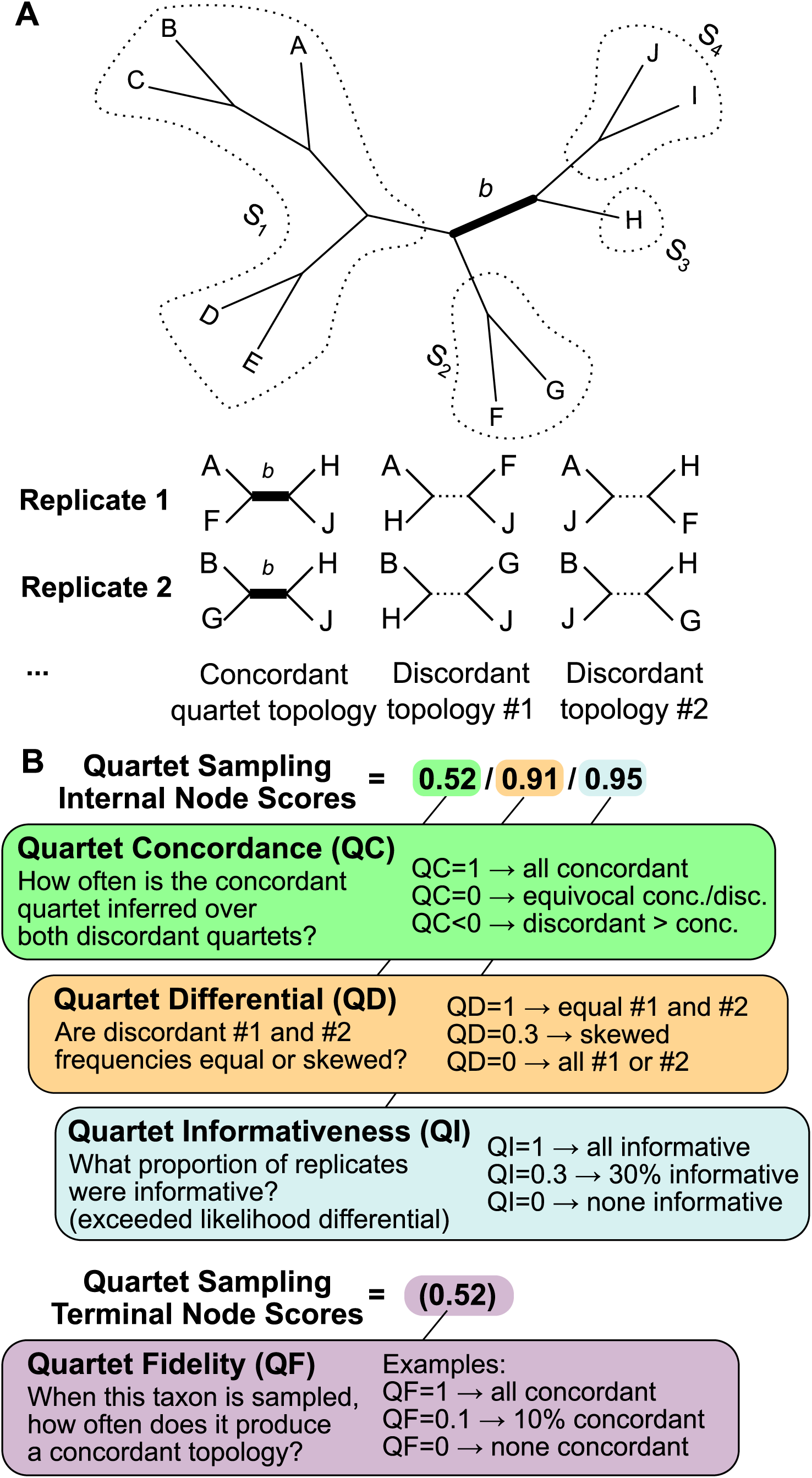
Description of the Quartet Sampling method. (A) The focal branch “*b*” divides the phylogeny into four subclades {*S*_1_*, S*_2_*, S*_3_*, S*_4_} from which tips (A–J) are sampled. Two replicates with different sampled tips for the given branch are shown with the three possible unrooted topologies (one concordant and two discordant). (B) Each internal branch is labeled with a set of three scores (QC/QD/QI), which offer different, but complementary, information. Terminal branches are evaluated by the QF score, which reports the frequency of a taxon generating concordant topologies. (See Materials and Methods for full details and Supplementary Methods for a technical description.)

**Table 1:** Quartet Sampling (QS) score interpretation. See text for details.

| Example QS Score<br>(QC/QD/QI)* | Interpretation |
| --- | --- |
| 1.0/-/1.0 | **Full support:** All sampled quartet replicates support the focal branch (QC=1) with all trees informative when likelihood cutoffs are used (QI=1). |
| 0.5/0.98/0.97 | **Strong support:** A strong majority of quartets support the focal branch (QC=0.5) and the low skew in discordant frequencies (QD≈1) indicate no alternative history is favored. |
| 0.7/0.1/0.97 | **Strong support with discordant skew:** A strong majority of quartets support the focal branch (QC=0.7), but the skew in discordance (QD=0.1) indicates the possible presence of a supported secondary evolutionary history. |
| 0.05/0.96/0.97 | **Weak support:** Only a weak majority of quartets support the focal branch (QC=0.05), and the frequency of all three possible topologies are similar (QD≈1). |
| 0.1/0.1/0.97 | **Weak support with discordant skew:** Only a weak majority of quartets support the focal branch (QC=0.1), and the skew in discordance (QD=0.1) indicates the possible presence of a supported secondary evolutionary history. |
| -0.5/0.1/0.93 | **Counter-support:** A strong majority of quartets support one of the alternative discordant quartet arrangements history (QC<0; QD expected to be low). |
| 1/0.97/0.05 | **Poorly informed:** Despite supportive QC/QD values, only 5% of quartets passed the likelihood cutoff (QI=0.05), likely indicating few informative sites. |
| 0.0/0.0/1.0 | **Perfectly conflicted:** The (unlikely) case where the frequency of all three possible trees was equal with all trees informative, indicating a rapid radiation or highly complex conflict. |

Notes: \* QC = Quartet Concordance; QD = Quartet Differential; QI = Quartet Informativeness.

In this study, we (1) describe the features, parameters, and interpretation of the QS method, (2) validate the QS method with simulations, and (3) apply the QS method to recently published large-sparse and phylogenomic datasets at timescales spanning from Viridiplantae to sub-generic clades. We demonstrate that the QS method is a flexible and computationally tractable method for examining conflict and support in large datasets. While not a panacea, we argue that the QS framework makes import steps in addressing many of the issues of branch support discussed above, and hope it encourages additional discussion, testing, and innovation of new phylogenetic evaluation methods. More broadly, the results presented herein contribute to the broader discussion about moving the plant tree of life beyond the goal of resolving a single, universal “Species Tree” (Hahn and Nakhleh, 2015; Smith et al., 2015), and into a future where we more fully explore and appreciate the complex “multiverse” of evolutionary histories manifest throughout the plant tree of life.

## MATERIALS AND METHODS

### Quartet Sampling

The Quartet Sampling (QS) procedure outlined here was inspired by aspects from several quartet-based and concordance methods, most particularly the process originally outlined by Hinchliff and Smith (2014b). The QS method takes an existing phylogenetic topology (which can be inferred by any method) and a molecular dataset (not necessarily the one that generated the phylogeny) and separately evaluates one or more internal branches on the given phylogeny. The QS method (Fig. 1) was designed to rapidly and simultaneously assess the confidence, consistency, and informativeness of internal tree relationships, and the reliability of each terminal branch.

For a given phylogeny, each observed internal tree branch partitions the tree into four non-overlapping subsets of taxa (Fig. 1A). These four sets of taxa (called a “meta-quartet” by Zhou et al., 2017) can exist in three possible relationships: the concordant relationship that matches the configuration in the given topology, and two alternative discordant configurations. The QS method repeatedly and randomly samples one taxon from each of the four subsets and then evaluates the likelihood all three possible phylogenies given the sequence data for the randomly selected quartet spanning that particular branch.

For each quartet sampled for the focal branch, the likelihood is evaluated (using the aligned sequence data) for all three possible topologies that these four sampled taxa can take (currently using RAxML or PAUP*, though other likelihood calculators could be substituted; Stamatakis, 2014; Swofford and Sullivan, 2003). The quartet topology with the best likelihood is then recorded and tabulated across all replicates. This process generates a set of counts (across all replicates per branch) where either the concordant or each of the two discordant relationships had the best likelihood. This procedure can be carried out by evaluating the likelihood of the complete alignment for each quartet (i.e., in a single-matrix framework) or by randomly sampling from individual gene/partition alignments from a multi-gene or genome-wide alignment (i.e., in a multi-gene tree coalescent framework).

Several refined options can be specified. For example, a minimum number of overlapping non-empty sites for all four taxa involved in a quartet can be specified to ensure calculations are performed on data rich subsets. Additionally, a parameter of a minimum likelihood differential may be set. If the most-likely topology (of the three) does not exceed the likelihood of the second-most-likely phylogeny by the set threshold, then the quartet is considered “uninformative” and tabulated separately. In summary, the QS method generates counts of the three possible topologies (and uninformative replicates) for each internal branch by sampling replicates using unique quartets of taxa spanning the particular branch.

The QS method uses these resampled quartet tree counts to calculate three scores for each internal branch of the focal tree (Fig. 1B, Table 1, and Appendix S1; see Supplemental Data with this article). The QC (Quartet Concordance) score is an entropy-like measure (similar to the ICA score; Salichos et al. 2014) that quantifies the relative support among the three possible resolutions of four taxa. When the most commonly sampled topology is concordant with the input tree, then QC takes positive values in the range (0,1]. Thus, QC equals 1 when all quartet trees are concordant with the focal branch. When one of the discordant topologies is the most commonly resampled quartet, QC takes negative values in the range [–1,0), approaching –1 when all quartet trees are one of the two discordant phylogenies. When support is evenly split among the three alternative topologies (or two if only two of the three possible are registered as having an optimal likelihood across all replicates), QC equals 0.

The QD (Quartet Differential) score uses the logic of the *f* - and *D*-statistics for introgression (Reich et al., 2009; Green et al., 2010; Durand et al., 2011; Pease and Hahn, 2015) and measures the disparity between the sampled proportions of the two discordant topologies (though with quartet topology proportions, rather than site frequencies). The QD score does not specifically quantify introgression nor identify introgressing taxa, but does indicate that one alternative relationship is sampled more often than the other. Low values of QD indicate that there is one preferred topology among the two discordant topologies, a potential indication on the given branch of a biased biological process beyond background lineage sorting, including confounding variables such as introgression, strong rate heterogeneity, heterogeneous base compositions, etc. QD varies in the range [0,1] with a value of 1 meaning no skew in the proportions of the two discordant trees and the extreme value of 0 meaning that all discordant trees sampled are only from one of the two possible alternative relationships.

The QI score (Quartet Informativeness) quantifies for a given branch the proportion of replicates where the best-likelihood quartet tree has a likelihood value that exceeds the quartet tree with second-best likelihood value by a given differential cutoff. This ensures that replicates are not counted as being concordant or discordant when the molecular data are effectively equivocal on the topology (i.e., when two of the three possible quartet topologies have nearly indistinguishable likelihood scores). QI is measured in the range [0,1], which indicates the proportion of sampled quartets that exceeded the cutoff. A QI value of 1 means all quartets are informative, while a value of 0 indicates all quartets were uncertain (i.e., no significant information for the given branch). The QI measure of branch informativeness works in conjunction with QC and QD to distinguish between branches that have low information versus those with conflicting information (i.e., high discordance).

Finally, for each terminal taxon, a QF (Quartet Fidelity) score is calculated to report the proportion of total replicates (across all branches tested) where the given taxon was included in a quartet resulted in a concordant quartet topology. QF is therefore similar in approach to a “rogue taxon” test (Wilkinson, 1996; Aberer et al., 2012). However, an important distinction is that RogueNaRok (Aberer et al., 2012) uses taxonomically complete bootstrap replicates to compute these scores rather than resampled subtrees, and thus are subject to the same issues as bootstrap scores themselves in phylogenomic analyses (i.e., RogueNaRok will not report rogue taxa when all bootstrap scores are 100). For a given taxon, the QF score is measured in the range [0,1] as the proportion of quartet topologies involving the taxon that are concordant with the focal tree branch. Therefore, a QF value of 1 indicates a given taxon always produces concordant topologies across all internal branches where it was sampled for in a quartet. QF values approaching zero indicate mostly discordant topologies involving this taxon, and may indicate poor sequence quality or identity, a lineage-specific process that is distorting the phylogeny, or that the taxon is significantly misplaced in the given tree. Note that QF differs specifically from QC, QD, and QI by being a taxon-specific test across internal branch tests rather than an internal branch-specific test.

Collectively, these four tests represent a means to distinguish the consistency of a branch (QC), the presence of a secondary evolutionary history (QD), the amount of information regarding a branch (QI), and the reliability of individual taxa in the tree (QF; Fig. 1B and see Table 1). Therefore, QS tests disentangle these effects rather than have them conflated under a summary score as in standard measures of phylogenetic support. A full technical description of the QS method is included in Appendix S1.

### Implementation of QS

We implemented the above procedure in a Python-based program called *quartetsampling*, which samples an alignment randomly to generate many representative quartet topology replicates for each internal branch in a corresponding focal tree (<u>https://github.com/fephyfofum/quartetsampling</u>). This procedure has a number of advantages over NBS for larger datasets. First, unlike NBS and RBS, alignment columns are not resampled, which allows sparse alignments to be used. Second, the number of likelihood calculations that are required is the number of internal branches in the tree multiplied by the number of replicates per branch multiplied by three possible topologies. Since computation time scales linearly with the number of taxa, individual replicates are fast, and the computations can be readily parallelized across processors and furthermore discretized across systems (with results combined later). This allows QS to be efficiently applied to large alignments beyond the practical limits of NBS and PP. The most extensive computational time was for the Zanne et al. (2014b) 31,749 taxon dataset (see below), which we ran on the Wake Forest University DEAC high-performance cluster using 8 nodes with 16 CPU each. This analysis completed 200 replicates for the full tree in 13 hours. Smaller genome-wide datasets finished 1000 gene-tree replicates on quad-core desktops approximately 12 hours. The conventional multi-gene datasets took only a few minutes to a few hours to run on a standard desktop.

Although the Shimodaira-Hasegawa-like approximate likelihood ratio test (SH-aLRT; Guindon et al., 2010) was by far the fastest method we consider here, the QS was fast enough for large scale analyses. QS can also be applied separately to individual focal branches, allowing for more thorough exploration of particular branches of interest. Furthermore, the QS does not require the tree tested to be the maximum likelihood topology, a requirement for SH-aLRT. For our simulated data, we found that performing 200 QS replicates per branch was adequate to achieve low variance in QS score (Fig. 2A). As would be expected, more replicates per branch should generally be used for larger trees to sample a greater fraction of the total possible quartets.

**Fig. 2.**
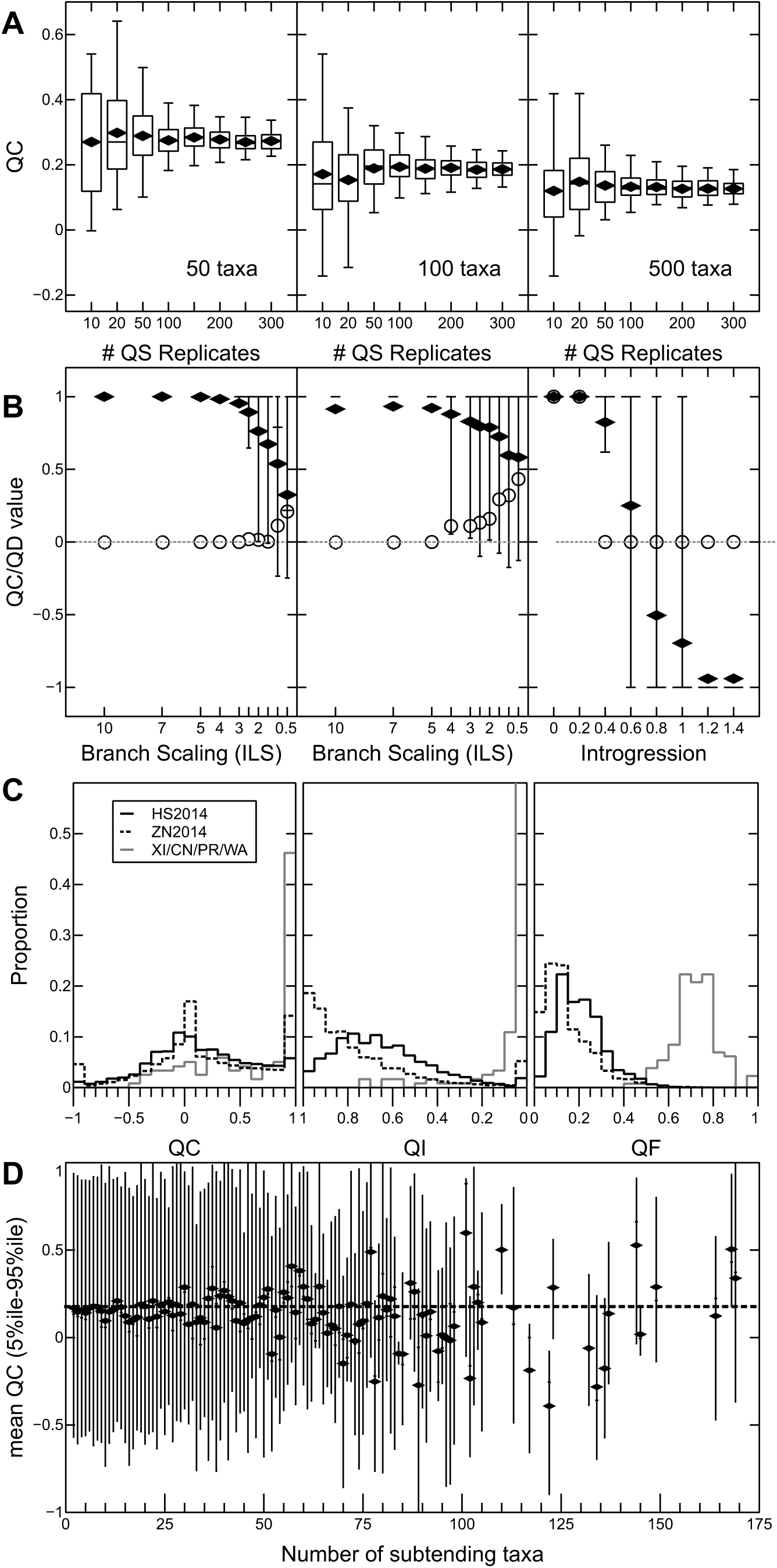
Results of Simulation Testing of the Quartet Sampling Method. (A) QC values converge on a central value with increasing numbers of replicates from randomly selected branches from simulated trees with 50, 100, and 500 taxa. (B) Mean QC (solid diamond) and QD (open circle) values with 5%ile to 95%ile (whiskers) across 100 replicates for branches QS16 (left) and QS12 (middle) for a simulated tree (Appendix S5) where the tree branch lengths were scaled by the factors on the *x*-axis (i.e., 1 is the original tree). As expected, shorter branch lengths will increase the level of incomplete lineage sorting (ILS) and thus lower the QC scores. The right panel shows branch QS11 from the simulated tree with increasing levels of introgression introduced by simulation. As expected, QC and QD values decrease with increasing introgression. (C) Distributions of QC, QI, and QF values for HS2014 (black), ZN2014 (dotted black), and XI2014/CN2015/PR2016/WA2017 (similar distributions; gray solid). (D) Mean QC values (diamond) with 5%ile to 95%ile (whiskers) for branches in HS2015 binned by the number of subtending taxa (i.e., moving root-ward in the tree left-to-right). Overall mean is shown with horizontal dotted line.

Furthermore, some branches, especially in large trees, may be entirely unsupported by the alignment due to a lack of sampling overlap among appropriate taxa (i.e., no sites in the alignment contain data from each of the four subsets of taxa; Fig. 1A). Therefore, no phylogenetic information exists to inform the branch (i.e., they are not “decisive” *sensu* Steel and Sanderson, 2010). The QS procedure identifies these branches, rather than discarding them or ambiguously labeling them as having “low support.”

#### Guidelines for interpretation of QS support values

An important consideration with any measure used to ascertain confidence is precise interpretation. We provide a concise visual description of the tests (Fig. 1) and a table describing example scores and their interpretations (Table 1). Particularly notable is that the QS method not only can “support” or “fail to support” a given branch hypothesis, but also can offer “counter-support” for an alternative branch (as in the IC/ICA scores; Salichos et al., 2014; Kobert et al., 2016; Zhou et al., 2017). Therefore, even “inaccurate” branch hypotheses can offer information as “counter-support” for an alternative quartet topology (i.e., the degree of negativity of the QC score; for examples see Fig. 6).

The QS scores we have described calculate the sensitivity of the resolution of a particular branch to different combinations of taxa sampled around that branch. Each QS replicate calculates whether the four sampled taxa support the resolution of the branch found in the tree over the alternative resolutions. This framework is similar to the interpretation made by those using taxon jackknife analyses for outgroup sensitivity (e.g., Edwards et al., 2005) and the IC score when used with incomplete trees (Kobert et al., 2016; Zhou et al., 2017). We argue that this interpretation is richer in information than the NBS, and, in simulations, the QC score also appears to more conservatively and accurately assign high support values to branches that are present in the true tree (i.e., relatively low false positive rates, at least when the likelihood threshold is small, i.e., in the range of *∼*2 used here; Appendix S2). QC scores are particularly helpful for clarifying strength of support for branches with concordant tree frequencies not close to 1 (Appendix S3).

#### Generation and evaluation of simulated phylogenies

We first tested the method by generating simulated phylogenies under the pure birth (birth = 1) model of evolution with 50, 100, and 500 tips using `pxbdsim` from the `phyx` toolkit (Brown et al., 2017). Using these trees, we generated 1000 bp alignments (no indels) under the Jukes-Cantor model with INDELible v. 1.03 (Fletcher and Yang, 2009). Trees were scaled so that the average branch lengths were about 0.2, based on the observation that this generated reasonable trees with most branches recovered correctly from ML analyses. Using the same procedure, we also simulated trees with 500 tips and associated alignments with ten nucleotide partitions, each with 500 sites under the Jukes-Cantor model. We simulated both the full alignment with partitions and a modified randomly resampled sparse alignment to examine the behavior of QS in the presence of missing data (see Appendix S1 for details). These partitioned and sparse alignments had the same qualitative features as the full alignment.

Unlike the NBS method, which generates a set of trees from which branch support is estimated, the QS method requires only a single input topology for which branch support will be measured. We calculated QC, QD, QI, and QF scores for the true underlying tree as well as the ML tree generated by RAxML, but we focus on results for the ML tree. To examine how the number of replicates impacts the QS precision, we conducted simulations varying the number of replicates for randomly drawn branches in the simulated trees (Fig. 2A; Appendix S4). Based on these simulations, we elected to use 200 replicates per branch, since the variance in the QC score was generally low across all tree sizes when this many replicates were performed. We used RAxML and PAUP* to estimate the ML for the three alternative topologies for each QS replicate (using the `-f N` option and the GTRGAMMA model in RAxML). We also calculated branch-specific QC/QD/QI and taxon-specific QF scores using likelihood differential cutoffs of Δ*L* = 0 (no filtering) and Δ*L* = 2.0, which requires stronger conflicting signal to interpret branches in the input tree as unsupported.

Additionally, we generated a simulated 20-taxon tree using `pxbdsim` from `phyx` (Brown et al., 2017) with variable branch lengths (Appendix S5). For 100 replicates, we generated twenty 5 kb nucleotide sequences over this tree using `ms` (Hudson, 2002), inferred a concatenated tree using RAxML (Stamatakis, 2014), and used this inferred tree and simulated alignment as the inputs for QS. Population parameters were set at *µ* = 1*×*10^-8^ and *N*_*e*_ = 10^5^. To simulate increasing amounts of ILS, we shortened the times between speciation events by scaling all branch lengths by factors ranging from 0.5 to 10 and repeated these simulations (Fig. 2C). Additionally, using the original tree scaled by a factor of 2, we added introgression of varying intensity between “taxon 6” and “taxon 7” (using the migration parameter in `ms` from 0 to 1.4/4*N*_*e*_ migrants per generation). Additional details can be found in Appendix S1.

#### Testing of Empirical Datasets

We evaluated five recent large-scale phylogenies, including (1) a 103-transcriptome dataset spanning Viridiplantae from Wickett et al. (2014, abbreviated hereafter as “WI2014”), (2) two large-sparse phylogenies spanning land plants from Hinchliff and Smith (2014b, “HS2014”) and Zanne et al. (2014b, “ZN2014”), and (3) phylogenies spanning Magnoliophyta (angiosperms) with hundreds of genes from Xi et al. (2014a, “XI2014”) and Cannon et al. (2015b, “CN2015”). Additionally, to demonstrate the utility of this method at medium and short time scales, we evaluated two whole transcriptome datasets from the wild tomato clade *Solanum* sect. *Lycopersicon* from Pease et al. (2016b, “PE2016”) and carnivorous plants from the order Caryophyllales from Walker et al. (2017c, “WA2017”). Finally, we tested this method on a more typical medium-sized multi-locus dataset from Polypodopsida (ferns) from Pryer et al. (2016b, “PR2016”), such as might appear in many phylogenetic studies of large subgroups. Data for these studies were obtained from datadryad.org and iplant.org (Hinchliff and Smith, 2014a; Matasci et al., 2014; Xi et al., 2014b; Zanne et al., 2014a; Cannon et al., 2015a; Pease et al., 2016a; Pryer et al., 2016a; Walker et al., 2017b) (additional details and results in Appendix S1).

In addition, we analyzed the datasets using 200 individual gene trees for XI2014 and WA2017, and 1000 gene trees for PE2016 and WI2014. For these datasets, quartets are sampled as usual, but only the individual gene sequence alignments are assessed. These phylogenies were all evaluated using a minimum alignment overlap per quartet of 100 bp and a minimum likelihood differential of 2 (i.e., the optimal tree’s log-likelihood must exceed the second-most likely tree by a value of at least 2). We also calculated the phylogenies with and without partitioning in RAxML, but in all cases the partitioned datasets did not qualitatively differ from the results of the unpartitioned datasets. These data are provided as supplementary data, but are not shown here.

We also either re-calculated other measures of branch support or used values from the published studies for comparison to the QS method for each phylogeny, except HS2014 and ZN2014 where the size and sparseness of the datasets prohibited the calculation of other measures of support. For the datasets from CN2015, PR2016, WA2017, and XI2014 100 replicates each of RAxML NBS and SH-test were performed. Additionally, PP scores for PR2016 were calculated using MrBayes (Ronquist et al., 2012), and IC scores for calculated for Walker et al. (2017c). For PE2016 and WI2014, RAxML NBS, MP-EST, or IC scores were taken from published values. Finally, we also calculated QF scores and rogue taxon scores using RogueNaRok (Aberer et al., 2012) to compare these two measures, particular for the large-sparse ZN2014 dataset (for details and results, see Appendix S1). Data and results from the simulations and empirical studies are available at Dryad (http://dx.doi.org/10.5061/dryad.6m20j).

### RESULTS AND DISCUSSION

#### Simulation analyses

We tested the consistency and reliability of QS on a set of simulated phylogenies. The QC scores converge (with decreasing variance as expected) on a consistent mean value for each branch as the number of replicates increased (Fig. 2A). Sampling 200 quartets per branch reduced the variance to less than 0.003 in all cases, and can be seen as a generally a reasonable number of replicates. As these are branch-specific tests, branches of interest can be tested individually at much higher numbers of replicates without the need to re-test the entire tree. Additionally, we simulated sequences over a standard phylogeny (Appendix S5), then simulated increasing ILS by shortening branch lengths and introgression via migration. As expected, QC scores that measured concordance decreased in both cases due to the increased presence of discordant sites and QD scores that measure skew in discordance decreased dramatically with increasing directional introgression (Fig. 2B). We also found that while QC and QD both measure discordance levels, they are not strictly correlated measures. As QC goes to the limits of its range [–1,1], QD values tend to have more extreme values that were due to a lack of discordant trees (QC near 1) or high frequency of one discordant tree (QC near –1). Applying a minimum log-likelihood differential threshold to small trees tended to push scores toward extremes, resulting in more 0s and 1s (Appendix S2). Finally, we found that those datasets with lower QF score generally identified more rogue taxa than inferred by RogueNaRok, despite the different data inputs and analysis frameworks (Appendix S1).

#### QS analyses of major land plant lineages

The primary goal of this study was to use QS to reanalyze and compare several recent speciose and phylogenomic datasets to address ongoing debates of phylogenetic relationships in the plant tree of life. We used QS methods to evaluate two of the most speciose phylogenies of land plants currently available from Hinchliff and Smith (2014b, Fig. 3) and Zanne et al. (2014b, Fig. 4), and one of the most comprehensive phylogenies of Viridiplantae from Wickett et al. (2014, Fig. 5). QS analyses were able to provide a broad scale summary of the stability of the datasets.

**Fig. 3.**
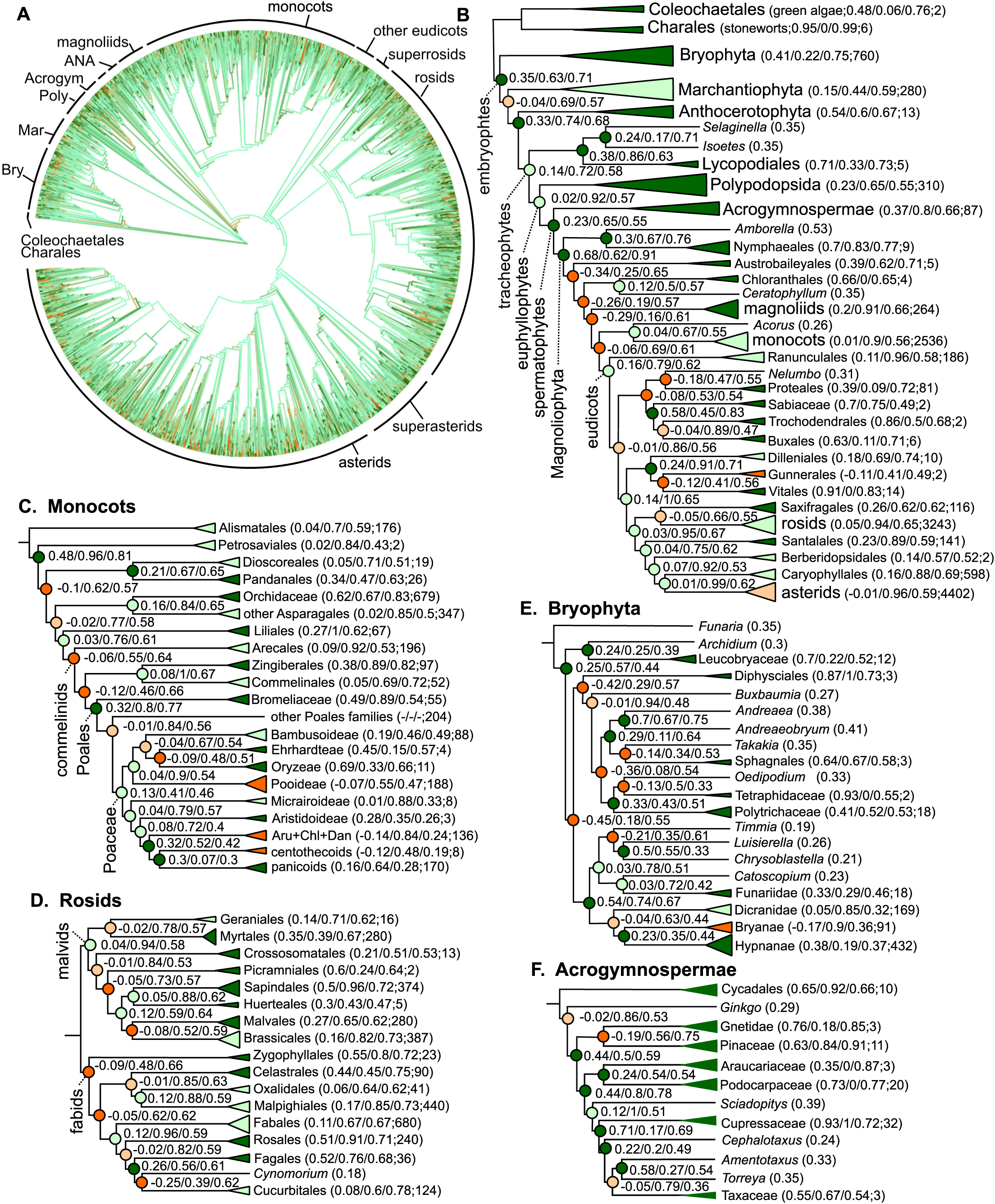
Phylogeny from Hinchliff and Smith (2014b). (A) Full phylogeny with heat map coloration of branches by QC scores for internal branches: dark green (QC>0.2), light green (0.2≤QC>0), light orange (0≤QC≤–0.05, or dark orange (QC>–0.05). (B) QC/QD/QI scores (200 replicates of full alignment) for major plant groups and key orders within angiosperms. QC/QD/QI scores after group names are for the ancestral branch (i.e., the “stem” branch), and a single QF score is shown for monotypic tips. Major subgroups groups are highlighted with vertical labels. (C) QS scores for monocots (excluding *Acorus*). (D,E,F) QS scores for rosids, Bryophyta, and gymnosperms. Abbreviations: Acro, Acrogymnospermae; ANA, ANA grade; Aru, Arundinoideae; Bry, Bryophyta, Chl, Chloridoideae; Dan, Danthonioideae; Mar, Marchantiophyta; Poly, Polypodopsida.

**Fig. 4.**
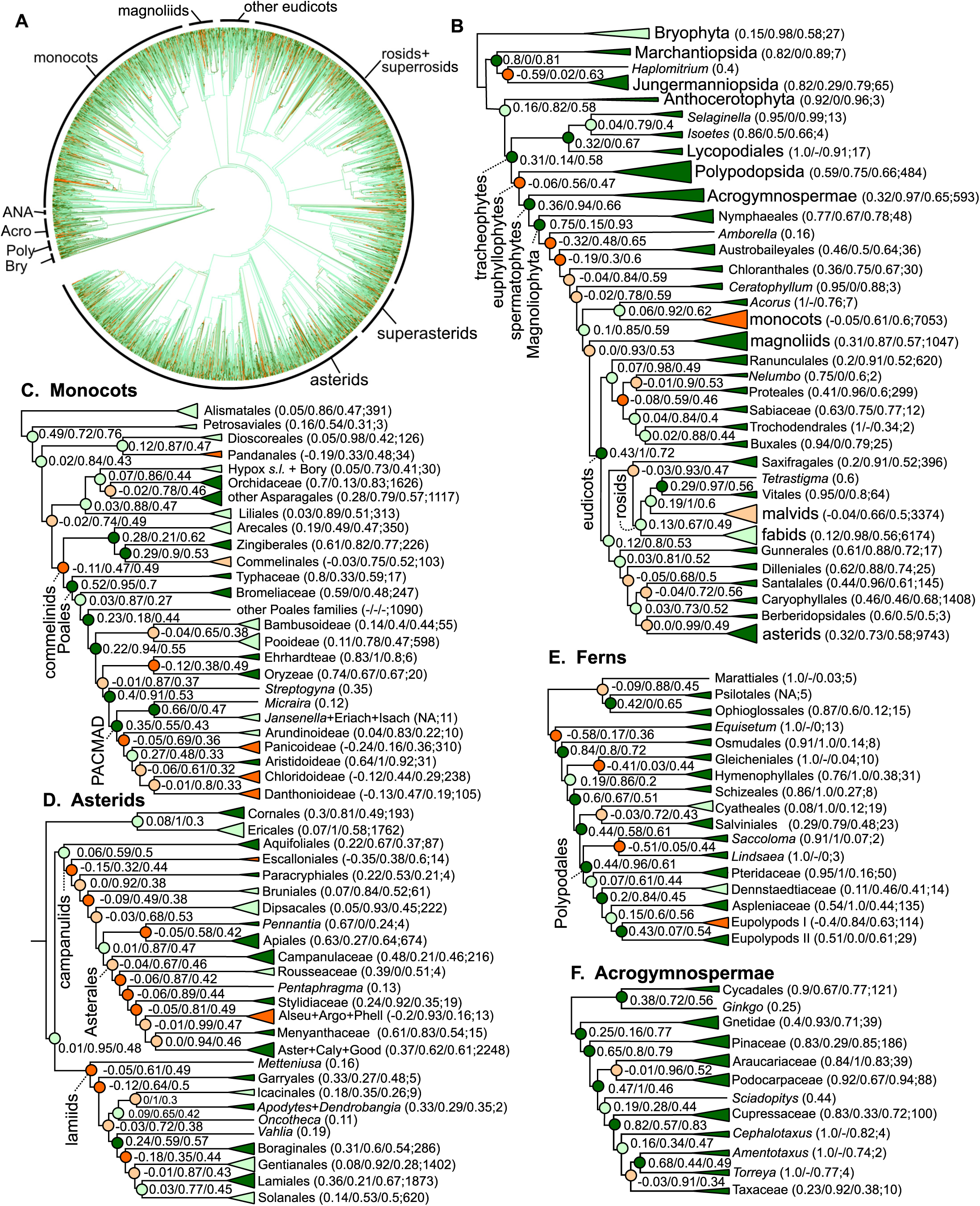
Phylogeny from Zanne et al. (2014b). (A) Full phylogeny with heat map coloration of branches by QC scores for internal branches using same color scheme as (Fig. 3). (B) QC/QD/QI scores (200 replicates of full alignment) for major plant groups and key orders within angiosperms, using same color scheme as (Fig. 3). (C) QS scores shown for monocots (except *Acorus*). (D) QS scores for asterids. (E) QS scores for fern lineages and (F) QS scores for gymnosperm lineages respectively. Abbreviations: Alseu, Alseuosmiaceae; ANA, ANA grade angiosperms; Argo, Argophyllaceae; Aster, Asteraceae; Bory, Boryaceae; Caly, Calycanthaceae; Eriach, Eriachneae; Good, Goodeniaceae; gym, gymnosperms; Hypox, Hypoxidaceae; Isach, Isachneae; Phell, Phellinaceae; Poly, Polypodopsida.

**Fig. 5.**
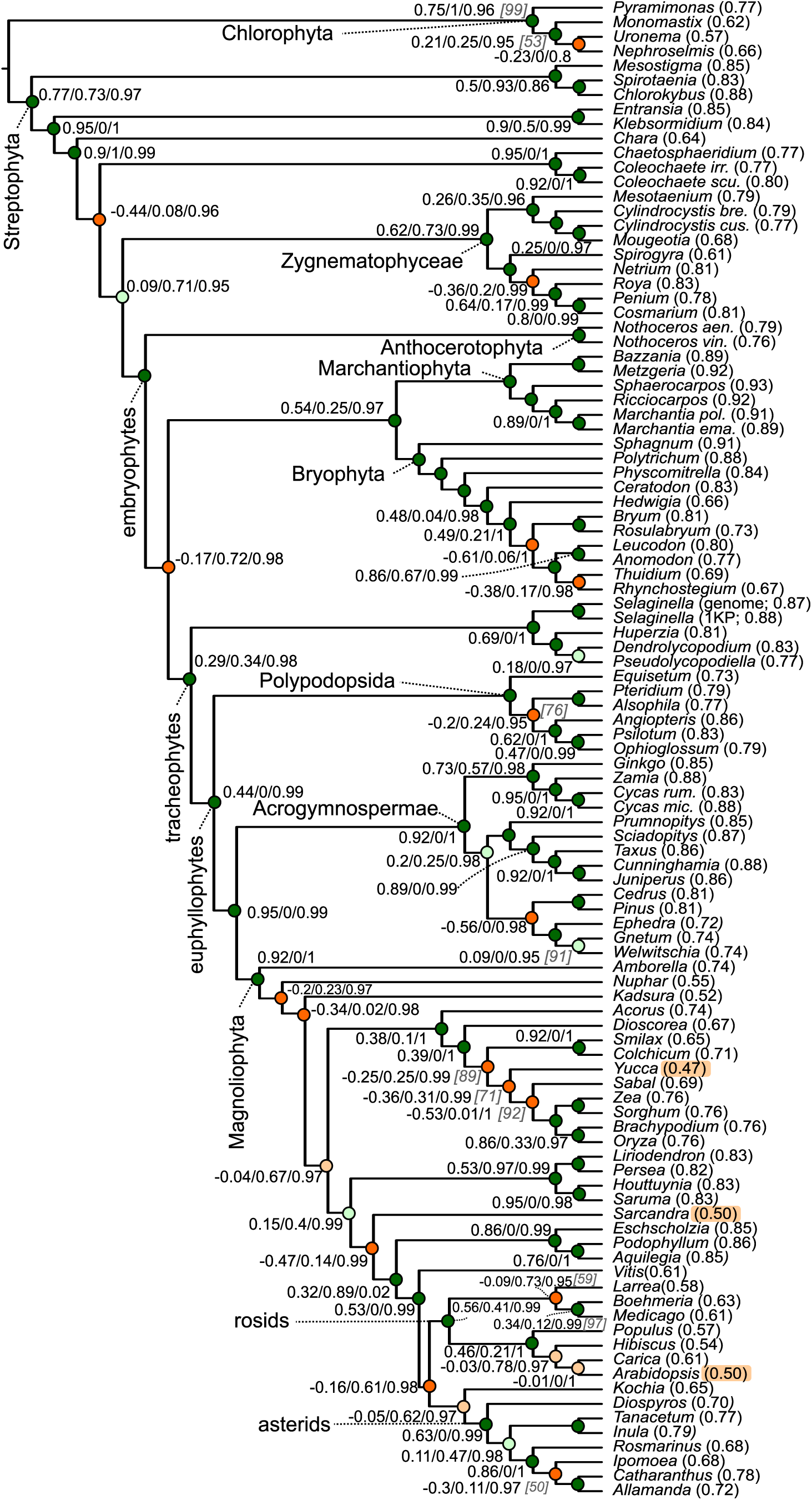
Maximum likelihood phylogeny spanning Viridiplantae from Fig. 2 in Wickett et al. (2014) with QC/QD/QI scores for 200 replicates of the full alignment. Nodes are colored according to QC score using same color scheme as (Fig. 3). Bootstrap values (italicized in square brackets) from Wickett et al. (2014) are shown for comparison. Missing QS or bootstrap values indicate a perfect score. The three taxa with the lowest QF values are highlighted. Species names have been excluded or abbreviated in the case where two congeners are included.

As expected, given the sparsity of the matrices for HS2014 and ZN2014 (96% and 82% missing characters, respectively), the proportion of informative quartets was low in both cases (mean QI of 0.15 and 0.35, respectively). Overall, the mean QC for the HS2014 (0.15; interquartile range (IQR) = [–0.13, 0.46]) and ZN2014 (0.17; IQR = [–0.10, 0.63]) were low compared to the less speciose phylogenies (Fig. 2C; Appendix S6). Notably, we found 33.4% and 29.8% of branches in HS2014 and ZN2014, respectively, had QC values less than –0.05 meaning that about a third of the branches in these consensus phylogenies reported not just “low support” for the given branch, but went further to report “counter-support” (i.e., a negative QC score) for one of the two alternative topological arrangements at that branch. Most major plant groups showed strong support in HS2014 and ZN2014 and all major groups showed strong support in WI2014 (Table 2). In contrast to strong support for major groups themselves, we found low support along the “backbone” relating these groups, in a manner consistent with most previous phylogenies of land plants.

**Table 2:** Quartet Sampling scores for key branches in the plant tree of life

| Branch | QC |  |  |  | Consensus Interpretation |
| --- | --- | --- | --- | --- | --- |
|  | HS2014 | ZN2014 | WI2014 | XI2014 |  |
| embrophytes (land plants) | 0.35 | n.t. | 1.0 | n.t. | strong support |
| tracheophytes (vascular plants) | 0.14 | 0.31 | 0.29 | n.t. | moderate-strong support |
| euphyllophytes (ferns + seed plants) | 0.02 | −0.06 | 0.44 | n.t. | low/variable support |
| spermatophytes (seed plants) | 0.23 | 0.36 | 0.95 | n.t. | strong support |
| Acrogymnospermae (gymnosperms) | 0.37 | 0.32 | 0.92 | 1.0 | strong support |
| Anthocerotophyta (hornworts) | 0.54 | 0.94 | 1.0 | n.t. | strong support |
| Bryophyta (mosses) | 0.41 | 0.15 | 1.0 | n.t. | moderate-strong support |
| Lycopodiophyta | 0.38 | 0.32 | 0.89 | n.t. | strong support |
| Magnoliophyta (angiosperms) | 0.68 | 0.75 | 0.92 | 0.95 | strong support |
| Marchantiophyta (liverworts) | 0.15 | 0.8 | 1.0 | n.t. | moderate-strong support |
| Polypodopsida (ferns) | 0.23 | 0.46 | 1.0 | n.t. | moderate support |
| Chloranthaceae + core angiosperms | −0.26 | −0.04 | n.m. | n.t. | counter-supported |
| Chloranthaceae + eudicots | n.m. | n.m. | −0.47 | n.t. | counter-supported |
| magnoliids | 0.20 | 0.31 | 0.53 | 0.54 | strong support |
| eudicots | 0.16 | 0.43 | 0.32 | 0.71 | moderate-strong support |
| asterids | −0.01 | 0.32 | 0.63 | 0.0 | low/variable support |
| rosids | 0.05 | n.m. | 1.0 | 0.25 | low/variable support |
| monocots (including *Acorus*) | 0.04 | 0.06 | 0.38 | n.t. | low/variable support |
| monocots (excluding *Acorus*) | 0.01 | −0.05 | 0.39 | 0.76 | low/variable support |
| Liliales + commelinids | 0.03 | n.m. | n.t. | n.t. | low support |
| Liliales + Asparagales | n.m. | 0.03 | n.t. | n.t. | low support |

The relationships among Marchantiophyta, Bryophyta (mosses), Anthocerotophyta, lycophytes, and “euphyllophytes” (i.e., ferns and seed-bearing plants) has been a matter of ongoing debate (Shaw et al., 2011). HS2014 places mosses as sister to the remaining land plants, but indicated counter-support (negative QC=–0.04) for a branch defining a common ancestor of liverworts with all other land plants to the exclusion of mosses (Figs. 3B). This suggested that the most common quartet branch among the replicates was not the branch displayed in the published tree. By contrast WI2014 shows strong support (with a high QC=0.67) for a common ancestor of mosses and liverworts (Fig. 5). ZN2014 shows weak support (low positive QC=0.15) for the branch separating mosses and liverworts from the rest of land plants. Therefore, while the topology of HS2014 was consistent with the order of many previous phylogenies (Nickrent et al., 2000; Qiu et al., 2006; Chang and Graham, 2011), the QS results collectively supported the alternative configuration of mosses and liverworts as sister groups (Fig 6A; see also Renzaglia et al., 2000; Zhong et al., 2013a).

**Fig. 6.**
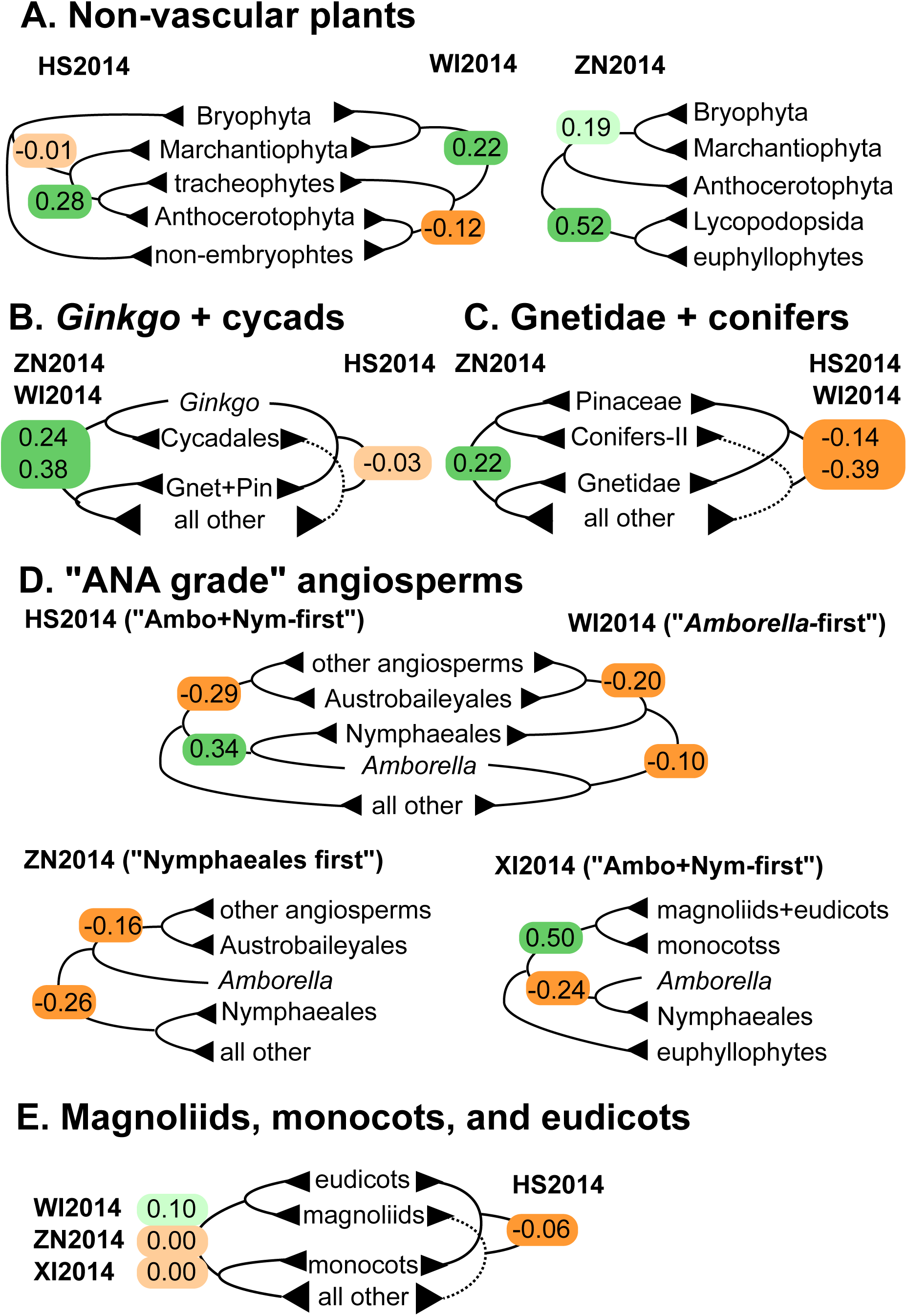
Key phylogenetic disagreements with QC scores using compared across various datasets. Branches for HS2014 and ZN2014 were resampled with 10000 replicates. Branches for WI2014 and XI2014 were exhaustively sampled (>1000 replicates). Highlighting on QC values follows the same colors as Fig. 3. “Conifers-II” refers to a hypothesized clade comprising the non-Pinales orders in Pinidae. Abbreviations: Gnet, Gnetidae; Pin, Pinidae.

In all three datasets, the monophyly of vascular plants was strongly maintained, even with the inclusion of *Selaginella* with its unusual GC content (Banks et al., 2011). The branch leading to *Selaginella* often had a lower QD value, possibly because of this biased composition, but a higher QF value, suggesting that it was not a misplaced (“rogue”) taxon. We also observed substantial discordance and counter-support for relationships tested among various bryophyte groups and key taxa in HS2014, possibly indicative of substantially under-appreciated hybridization among mosses (Nylander et al., 2004).

#### QS analyses of ferns

The branch establishing a “euphyllophyte” common ancestor of ferns and seed-bearing plants showed low QC scores and high QD scores in both HS2014 and ZN2014, indicating only a weak consensus but little indication of an alternative history (Table 2). Within ferns the arrangement of major clades in ZN2014 (Fig. 4E) was mostly consistent with the recently published phylogeny by The Pteridophyte Phylogeny Group (PPG I, 2016). Those clades whose relationships were counter-supported (Marratiales, Salviniales, Hymenophyllales) were discordant with the PPG-I consensus and other recent phylogenies (Pryer et al., 2004; Testo and Sundue, 2016) demonstrating the diagnostic utility of QS in highlighting suspect relationships. Some key areas of known high uncertainty (e.g., *Saccoloma*, *Lindsaea*, and *Equisetum*) were also highlighted with low or negative QC scores.

While QS was designed for large datasets, we also found that QS can perform well on smaller multi-gene datasets conventionally used for systematics studies. The QS scores for PR2016, with a 5778 bp alignment, were more conservative, but confirmed the conclusions of Pryer et al. (2016b) regarding the monophyly of maidenhair ferns (*Adiantum*) and its placement in a clade with the vittarioids. This analysis also revealed some counter-supported nodes (negative QC values) within the genus *Adiantum*.

#### QS analyses of gymnosperms

Another question that has attracted substantial historical debate is the relationships among the major gymnosperm lineages and angiosperms. Under QS evaluation, all four testable datasets indicated strong support for monophyly of gymnosperms (Table 2). However, the relationships among cone-bearing lineages differed among these four phylogenies. ZN2014 and WI2014 inferred a common ancestor of *Ginkgo* and cycads (consistent with Qiu et al., 2006; Bowe et al., 2000; Lee et al., 2011; Xi et al., 2013). While the HS2014 topology places cycads as sister to the remaining gymnosperms (i.e., not monophyletic with *Ginkgo*), the QS evaluation counter-supports this relationship. Therefore, even though HS2014 and WI2014 differed from ZN2014 in the topological relationship of these taxa, the QS analyses of these datasets indicated a consistent message of a *Ginkgo* and cycads common ancestor separate from the rest of gymnosperms (Fig. 6B).

This pattern of disagreeing topologies but consistent QS interpretation was observed again in the placement of Gnetales relative to the conifer linages (Fig. 6C). ZN2014 showed a common ancestor of Gnetales and Pinales (consistent with Lee et al. 2011). While a conflicting Gnetales and Pinaceae ancestor (distinct from other conifers) appeared in both HS2014 and WI2014 (i.e., the “Gnepine” hypothesis; Bowe et al., 2000; Xi et al., 2013), the negative-QC/low-QD scores in both cases (QC/QD=–0.19/0.56 and –0.67/0.0, respectively) indicate counter-support for a “Gnepine” ancestor and a strongly support alternative history. Collectively, these results suggests the monophyly of Pinales, but also offer some (albeit weak) evidence that warrants further examination of possible gene flow between Gnetales and Pinales.

#### QS analyses of ANA grade angiosperms

Few issues in angiosperm evolution have garnered more recent debate than the relationship among the so-called “ANA grade” angiosperms (Qiu et al., 1999), which include *Amborella*, Nymphaeales, and Austrobaileyales. Two questions surround the evolutionary history of the ANA grade angiosperms. First, what are the relationships among these lineages? Second, are the longstanding disagreements in inference of these relationships the result of genuine biological conflict (i.e., introgression, horizontal transfer, etc.), limitations in the data, or a methodological artifact (i.e., due to the depth of this branch, the monotypic status of *Amborella*, and/or the rapidity of the angiosperm radiation)?

On the first question, QS analyses of the datasets here lack support for “Nymphaeales-first” but finds support for both *Amborella*+Nymphaeales and “*Amborella*-first” (as found also by The Amborella Genome Project, 2013). While the resolutions of consensus phylogenies differ between the four testable datasets (WI2014 with “*Amborella*-first” hypothesis, ZN2014 with “Nymphaeales-first”, and HS2014 and XI2014 with *Amborella*+Nymphaeales), the branches surrounding the ANA-grade were all counter-supported (QC< 0) and biased in their discordance (QD< 0.2; Fig. 6D). ZN2014 offers weak support for *Amborella*+Nymphaeales, while XI2014 counter-supports this relationship. If this question is to be resolved, our results indicate additional datasets and analyses will be required.

On the second question, our analyses show low QD values that suggest a conflicting phylogenetic history may be present. Other studies have found bryophyte mitochondrial sequences present in *Amborella* (Rice et al., 2013; Taylor et al., 2015), which establishes the potential for introgression in these lineages. Overall, (1) the intense efforts to address these relationships without a resulting broad community consensus, (2) evidence of long-range introgression, and (3) the QS results shown here together suggest that a greater understanding of ANA-grade evolution likely lies in an examination of complex evolutionary histories rather than in a continuation of the debate over appropriate sampling or models (see also discussion in Shen et al., 2017).

#### QS analyses of “core angiosperms”

The three “core angiosperm” lineages (eudicots, monocots, and magnoliids) have transformed the biosphere, and thus a better understanding of the timing and order of their origins is of key concern. Consensus topologies disagree between ZN2014, WI2014, and XI2014 (with magnoliid+eudicot clade Figs. 4B, 5, 6E, Appendix S7) and HS2014 (with eudicot+monocots Fig. 3B). However, the QS analyses of HS2014 showed counter-support of an exclusive common ancestor of eudicots and monocots, suggesting that, despite disagreement among topologies, QS scores support a common ancestor for magnoliids and eudicots to the exclusion of monocots. Additionally, the placement of Chloranthaceae seems inextricably linked with the relationships of the three core angiosperm groups (see discussion in Eklund et al., 2004). However, the placement of this family remains unresolved by QS, since all tested configurations showed negative QC-value counter-support (Table 2).

#### QS analyses of monocots

In general, the arrangement of monocot orders in both HS2014 (Fig. 3C) and ZN2014 (Fig. 4C) agreed with recent consensus phylogenies (Givnish et al., 2010; Barrett et al., 2015; Givnish et al., 2016; McKain et al., 2016). Two exceptions are the placement of Liliales (Table 2), and general inconsistency of commelinid orders. From the QS results, we would cautiously infer that (1) the relationships among the commelinids are still unknown, (2) there may be uncharacterized secondary evolutionary history distorting the phylogenetic placement of these groups, and (3) likely the variable data from both Liliales and Arecales together have a joint effect that is causing inconsistency in the phylogenetic inference.

In Poaceae, QS analyses highlight the well-characterized discordance and complex relationships (e.g., Washburn et al., 2015; McKain et al., 2016). Even if someone were completely unfamiliar with the known controversies in monocots, QS scores would make abundantly clear this area of the phylogeny had highly conflicted data. The “BOP” clade itself and many clades with in the “PACMAD” clade were counter-supported by negative QC values in HS2014 and ZN2014. However, low QI values were observed in both HS2014 and ZN2014 for this clade, indicating that both datasets contain poor information. Therefore, QS serves as an effective diagnostic tool for identifying conflicted portions of larger phylogenies.

#### QS analyses of non-rosid/asterid eudicots

QS analyses are capable of identifying conflict and discordance due to rapid radiations. This is demonstrated well for the relationships among the superasterid groups (Caryophyllales, Berberidopsidales, Santalales, and asterids). A common pattern was found in HS2014, WI2014, XI2014, and ZN2014 of near-zero QC values (–0.03 to 0.08) that indicate weak consensus for the given relationships, strong QD values (0.97–1) that indicate no strongly competing alternative history, and low QI values (0.14–0.51) that indicate low information for branches. This led to a consensus QS interpretation of simple poor phylogenetic information, likely as a result of the rapid radiation of these lineages. Generally, these phylogenies tended to support weakly the controversial placement of Caryophyllales as most closely related to the eudicot ancestor.

#### QS analyses of rosids and asterids

Analysis of the rosids confirms that the QS method is capable of identifying rogue taxa. The QS scores identified a poorly supported relationship in HS2014 between *Cynomorium* and Cucurbitales (QC=–0.31). *Cynomorium*, a non-photosynthetic parasitic plant with unusual morphology, has been placed tenuously and variably in groups as diverse as Rosales (Zhang et al., 2009) and Saxifragales (Nickrent et al., 2005), so its poor score here was expected. This “rogue” status was corroborated by a below-average QF score of QF=0.18 (mean 0.21 for HS2014). This means that for quartets that include *Cynomorium* as a randomly sampled taxon, only 18% produced a quartet topology concordant with the HS2014 tree.

Published phylogenies of asterids indicate disagreement and substantial discordance (Soltis et al., 2011; Beaulieu et al., 2013; Refulio-Rodriguez and Olmstead, 2014). QS scores from ZN2014 supported the unusual hypothesis of a common Ericales+Cornales ancestor, weakly support the campanulid clade, and counter-support a common lamiid ancestor. The arrangement of families within Asterales either roughly conforms to Soltis et al. (2011) and Beaulieu et al. (2013), or counter-supports branches (QC< 0) that do not agree with these consensus phylogenies. However, most of the branches that define the relationships among asterid orders in ZN2014 were counter-supported by the data, though most have QC and QD values close to zero. This indicates a scenario of a rapid radiation rather than hybridization (though these are not mutually exclusive).

#### QS of shallow-timescale phylotranscriptomic datasets

So far, we have demonstrated the utility of quartet sampling on large, sparse, and conventional multi-gene alignments, which are often computationally intractable with other support measures. We have also shown for WI2014 that a relatively large and full occupied matrix from deep-timescale transcriptomic data can also be evaluated by QS. However, the QS method also can be used to rapidly evaluate phylogenetic support on genome-wide datasets with little missing data for shorter evolutionary timescales. We tested the QS method on two phylotranscriptomic datasets for the wild and domesticated tomato clade *Solanum* sect. *Lycopersicon* (Fig. 8A; Pease et al., 2016b) and carnivorous plants spanning the Caryophyllales (Fig. 8B; Walker et al., 2017c).

**Fig. 7.**
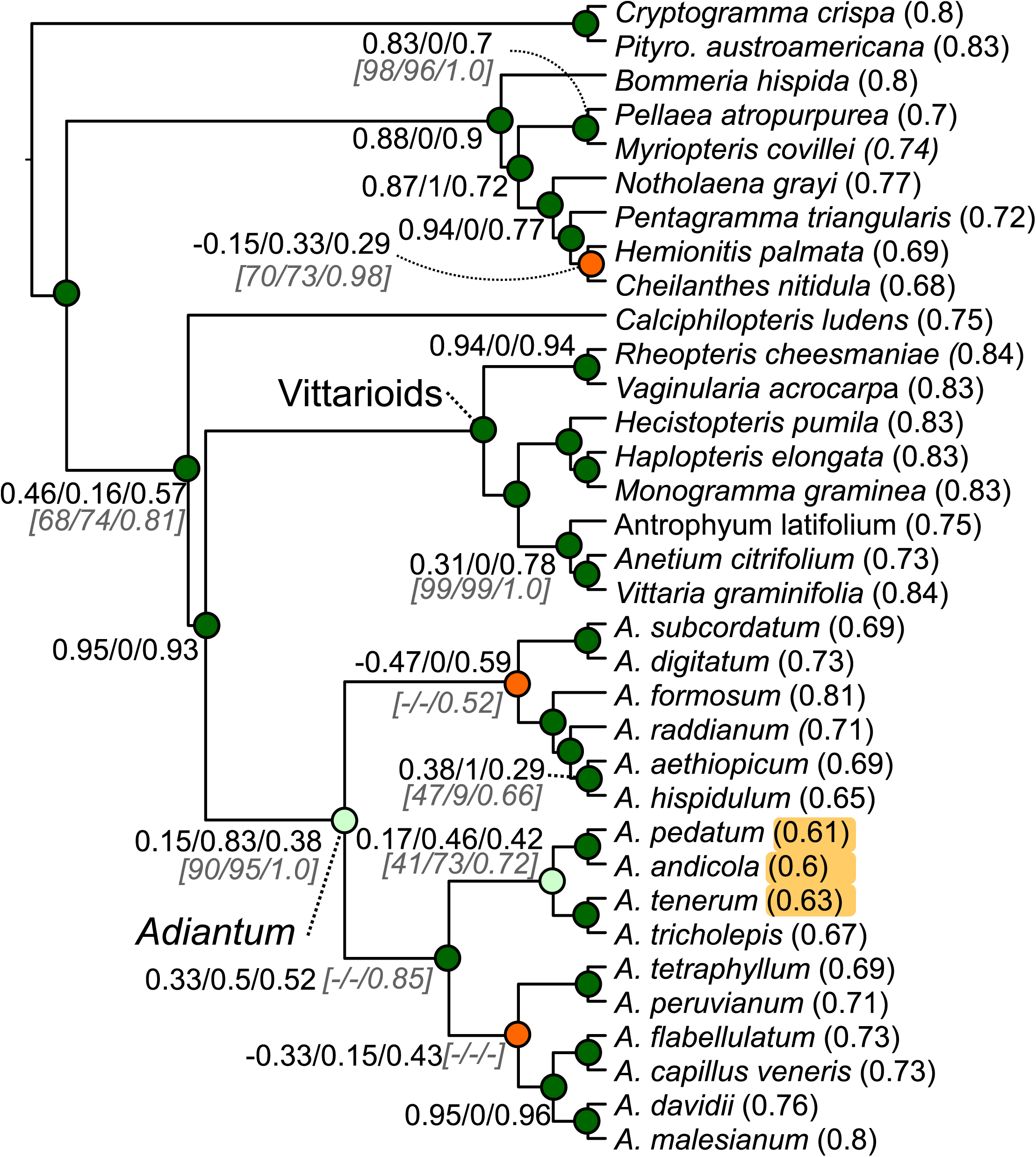
Phylogeny of Pteridaceae ferns from Pryer et al. (2016b) with QC/QD/QI scores for 200 replicates of the full alignment. Nodes are colored according to QC score using same color scheme as (Fig. 3). Bootstrap/SH-test/posterior probability values (italicized in square brackets) are shown for comparison. Omitted values indicate a perfect score. The three taxa with the lowest QF values are highlighted. Abbreviations: *Pityro*, *Pityrogramma*.

**Fig. 8.**
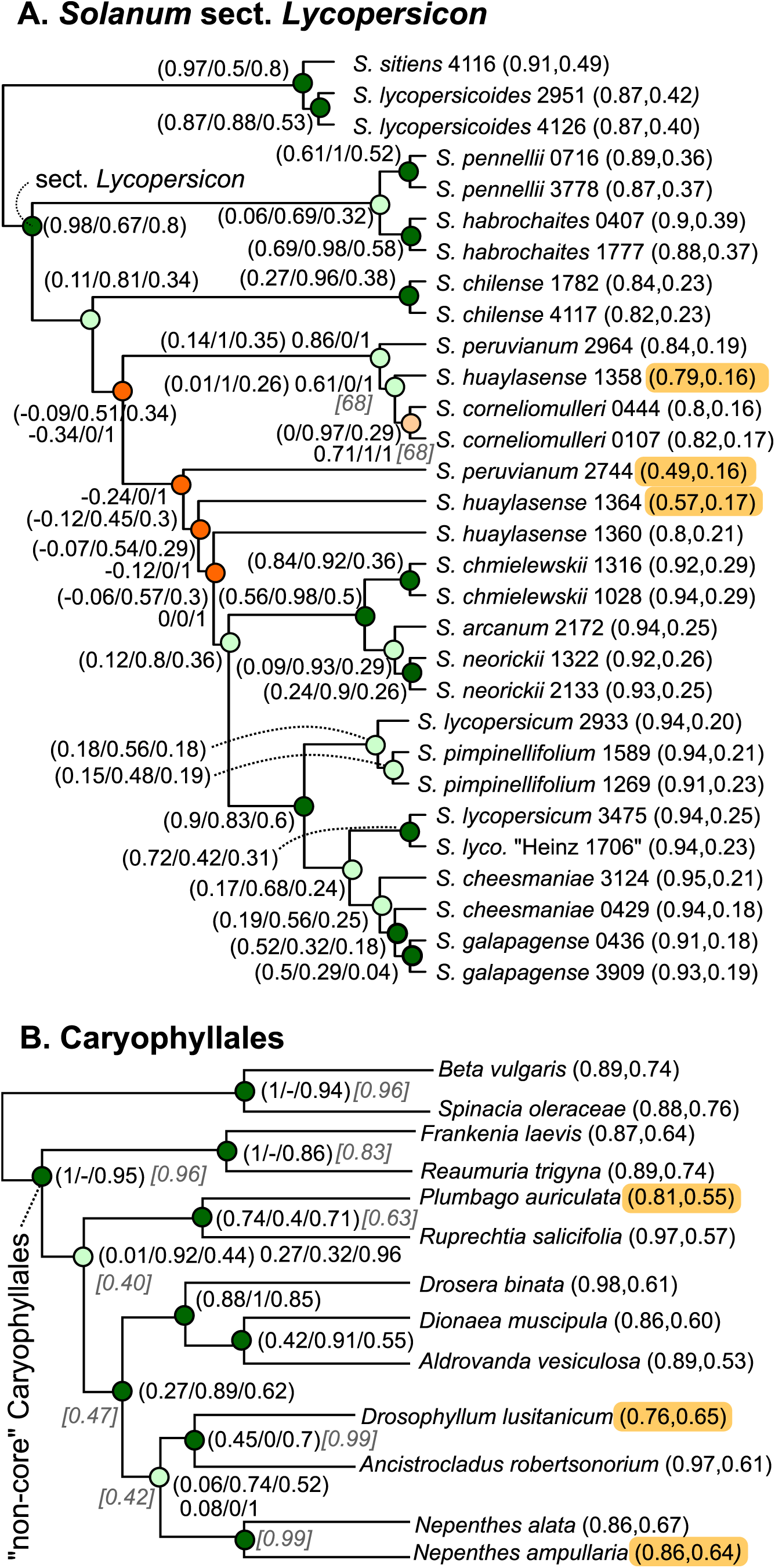
QS scores for phylogenies from whole-transcriptome data. Omitted values indicate a perfect score. Nodes are colored according to QC score using same color scheme as (Fig. 3). (A) Phylogeny of *Solanum* sect. *Lycopersicon* from Pease et al. (2016b) Bootstrap values (italicized in square brackets) are shown for comparison. (B) Phylogeny of Caryophyllales from Walker et al. (2017c) IC scores (light grey) are shown for comparison (all bootstrap and SH-test scores were 100). The three taxa with the lowest QF values are highlighted.

The *Solanum* phylogeny from Pease et al. (2016b) was inferred from the alignment of 33,105,168 nucleotide sites for 30 populations spanning all 13 wild and domesticated tomato species, and two outgroup species. As described in Pease et al. 2016b, this dataset contains a high level of phylogenetic discordance, but had a consensus phylogeny with 100% NBS support at all but two branches. However, gene tree analysis of this group showed evidence of massive phylogenetic discordance. When we applied QS to this phylogeny using the entire alignment, scores for many branches were also perfect (i.e., 1/–/1; Table 1). However, several of the other branches in the “Peruvianum group” species complex had lower QS scores in the full alignment (Fig. 8A). When gene trees were used (a random gene and quartet of taxa were chosen for 1000 QS replicates), all branches had QC< 1 in a manner consistent with the gene tree discordance found previously in this clade. We also observed the presence of low QD values within the major subgroups reported for this clade, indicating the presence of introgressive gene flow. In contrast, nodes defining the major subgroups showed high QC and QD values, indicate strong monophyly. This accurately captures the low discordance between groups versus high discordance within the major groups found by Pease et al. (2016b).

Most notably, the tree shown in Fig. 8A includes *S. huaylasense* accession LA1360. This accession has been known (both from Pease et al. (2016b) and other datasets) to mostly likely be a hybrid between populations from the green-fruited and red-fruited lineages (essentially those accessions above and below LA1360, respectively, in Fig. 8A). Thus, the inclusion of this putative hybrid lineage distorted the phylogeny as tree inference methods tried to cope with inherited and introgressed alleles from two separate groups to place this accession in a consensus location on the tree. While NBS scores were high for the branches surrounding the placement of LA1360, QS showed negative QC scores and low QD scores (QD=0 for full alignment). The low QD supports the presence of the alternative phylogenetic history that has been previously corroborated by other studies and the negative QC indicates counter support for the placement of this accession (see additional discussion in the Supplementary Results of Pease et al. 2016b). These data show that QS was able to distinguish between consistently supported relationships and branches known to have conflict due to introgression (whereas NBS does not).

An analysis of transcriptomes of carnivorous plants from Caryophyllales (Fig. 8B; Walker et al. 2017c) also highlighted the ability to dissect the dataset more effectively. The near-zero QC scores and low QD (0.32) scores for the ancestor of a clade containing *Plumbago* and *Nepenthes* for gene trees supported the hypothesis of Walker et al. (2017c) that introgressive gene flow may have occurred among these lineages. Evidence for placing *Drosophyllum* among the carnivorous Caryophyllales has been previously tenuous, and the QS analysis showed not only a low QF value of 0.76 (compared to the WA2017 mean QF of 0.89) for this taxon, but also low-QC/low-QD values for the two branches that form the clade with *Ancistrocladus* and *Nepenthes*. As with the tomato example above, this example demonstrates how QS scores can highlight an entire region that may be distorted by the inclusion of a taxon with a strong potential for a secondary evolutionary history (i.e., possible introgression).

#### Limitations and directions forward

Quartet Sampling is designed to efficiently evaluate phylogenetic information and to highlight conflict for one or more branches in a phylogeny. In the presentation here, QS is used to evaluate a single topology, and not for comparing alternatives topologies or performing any optimizations that might maximize QS scores. Therefore, QS does not suggest topological rearrangements and is purely evaluative. These and other directions should be explored in future studies as researchers develop more ways to examine uncertainty in large datasets.

Concurrently with our study, Zhou et al. (2017) have proposed the Q-IC method, a similar approach to QS. Both approaches use quartets to evaluate a focal tree. Both approaches can be used in a single-matrix or multi-gene tree framework, implemented in Q-IC by sampling from either a single tree distribution or from a gene tree set, and implemented in QS by analyzing either the whole alignment or by also randomly sampling individual gene-quartet combinations (as shown in Fig. 8). One key difference is that QS evaluates the relative likelihood of all three possible quartet configurations for each branch based on the alignment dataset, while Q-IC evaluates only the quartet topologies sampled from a dataset of topologies from “evaluation trees” (i.e., individual gene trees or a bootstrap/posterior distribution). These differences in data evaluation might make these approaches sensitive to different error types (e.g., gene tree topological estimation error versus likelihood estimation errors). Overall, we find these approaches complementary and their appropriateness dependent upon the data available and types question being asked.

### CONCLUSION

We reanalyzed several long-contested, key conflicts in the plant tree of life and describe a framework for distinguishing several causes of low phylogenetic branch support. For large datasets, traditional measures such as the bootstrap or posterior probabilities can be computationally intractable, may exhibit irregular behavior, or report high confidence despite substantial conflict. The QS framework provides a tractable means to analyze sparse datasets with tens of thousands of taxa but poor sequence overlap. QS provides a key function that has been missing from other support measures, namely the ability to distinguish among difference causes of low support that commonly occur in modern molecular phylogenies. We demonstrate this by reporting the existence of multiple conflicting but supported evolutionary histories at several key points in the plant tree of life (e.g., the placement of *Amborella*, possible widespread gene flow in the monocots, and notoriously difficult-to-place groups like *Cynomorium*). We hope that our discussions here will also lead to the development of other means for parsing the information contained within exponentially expanding molecular datasets. The artist Man Ray once remarked that “We have never attained the infinite variety and contradictions that exist in nature.” Overall, the picture painted by QS is one of substantial contradiction, but this conflict can be a richly informative (not just confounding) illustration of the interwoven evolutionary histories contained within the plant tree of life.

## ACKNOWLEDGEMENTS

The authors thank Ya Yang, Caroline Parins-Fukuchi, and Kathy Kron for helpful discussions, and Luke Harmon, Eric Roalson, Matt Pennell, Alexis Stamatakis, and an anonymous reviewer for valuable feedback on drafts. Computations were performed on the Wake Forest University DEAC Cluster, a centrally managed resource with support provided in part by the University.

## APPENDICES

### Appendix S1

Supplementary Methods providing a technical description of the QS method

### Appendix S2

Comparison of QC and bootstrap ICA (information criterion-all; Salichos, et al. 2014) scores on trees reconstructed from 100 simulated datasets with 50 taxa with 1,000 base pairs under a Jukes-Cantor model of evolution. Blue circles represent branches in the true tree, with the size of the circle proportional to the log of the number of substitutions. Red triangles represent branches not in the true tree.

### Appendix S3

Comparison of the rapid bootstrap and quartet sampling on the ML/PP consensus tree. For each branch, the RBS, QS (raw concordant frequency (Freq1), QC score), SH, and PP scores are presented (clockwise from top left in each legend). Black dots identify clades that are not in the true tree.

### Appendix S4

Shows the consistency of the frequency of concordant quartets (*f*_1_), QC, and QD toward a central value with increasing number of per-branch replicates for a randomly selected branch. Trees with 50 taxa (left), 100 taxa (center), and 500 taxa (right) are shown. Boxes show median ± IQR. Whiskers show 5^th^–95^th^ percentile, with values outside this range shown as circle points.

### Appendix S5

Simulated starting phylogeny used for the variation of simulated ILS and introgression levels shown in Fig. 2B.

### Appendix S6

Histograms (top row) showing the distributions of QC (left), QI (middle), and QF (right) values for the HS2014 dataset (green), ZN2014 (black), and smaller dataset (XI2014, CN2015, PR2016, WA2017) with similar distributions (orange). Scatter plots (bottom row) showing the close (but non-linear) relationship between QC and raw concordant quartet frequency (*f*_1_; left), bounded but otherwise uncorrelated relationships between QC and QD (middle), and QC and QI (right). See main text for dataset abbreviations.

### Appendix S7

Phylogeny of angiosperms from Xi et al. (2014a) with QC/QD/QI scores for 200 replicates of the full alignment and for 200 replicates from individual gene trees (in parentheses). Nodes are colored according to QC score using same color scheme as (Fig. 3). MrBayes PP/RAxML NBS values (italicized in square brackets) from Xi et al. (2013). are shown for comparison. Perfect scores for any given test are omitted or shown as ‘*’ indicates bootstrap of 100, while ‘-’ indicates a missing value. The three taxa with the lowest QF values are highlighted.

### Appendix S8

Phylogeny from Cannon et al. (2015b) with QC/QD/QI scores for 200 replicates of the full alignment. Nodes are colored according to QC score using same color scheme as (Fig. 3). Bootstrap values (italicized in square brackets) are shown for comparison. Perfect scores for any given test are omitted or shown as ‘*’ indicates bootstrap of 100, while “–” indicates a missing value. The three taxa with the lowest QF values are highlighted.

### Appendix S9

Relationship between QC and frequencies of the three possible alternative quartet topologies from QS runs on simulated data. Points represent branches in the trees, with the “test topology” axis identifying the frequency at which the topology consistent with the tree was recovered across all QS replicates for that branch, and the “alt n” axes identifying the frequencies of the two alternative (conflicting) topologies.

